# Anesthesia suppresses gastric myoelectric power in the ferret

**DOI:** 10.1101/2023.02.23.529745

**Authors:** Lorenzo Tomaselli, Michael Sciullo, Stephanie Fulton, Bill J. Yates, Lee E. Fisher, Valérie Ventura, Charles C. Horn

## Abstract

**Background:** Gastrointestinal myoelectric signals have been the focus of extensive research; although it is unclear how general anesthesia affects these signals, studies have often been conducted under general anesthesia. Here, we explore this issue directly by recording gastric myoelectric signals during awake and anesthetized states in the ferret and also explore the contribution of behavioral movement to observed changes in signal power.

**Methods:** Ferrets were surgically implanted with electrodes to record gastric myoelectric activity from the serosal surface of the stomach, and, following recovery, were tested in awake and isoflurane-anesthetized conditions. Video recordings were also analyzed during awake experiments to compare myoelectric activity during behavioral movement and rest.

**Key Results:** A significant decrease in gastric myoelectric signal power was detected under isoflurane anesthesia compared to the awake condition. Moreover, a detailed analysis of the awake recordings indicates that behavioral movement is associated with increased signal power compared to rest.

**Conclusions & Inferences:** These results suggest that both general anesthesia and behavioral movement can affect the amplitude of gastric myoelectric. In summary, caution should be taken in studying myoelectric data collected under anesthesia. Further, behavioral movement could have an important modulatory role on these signals, affecting their interpretation in clinical settings.

## 1. INTRODUCTION

Gastrointestinal (GI) myoelectric signals have been the focus of extensive research, especially regarding their relationship with gastric motility, but due to methodological constraints they currently have limited clinical use [1,2]. There is a critical need to determine the value of these signals for clinical application and to assess stability of these measures in different testing conditions. One significant testing variable has been the common use of general anesthesia during tests of myoelectric function in preclinical, as well as human studies [e.g., 3–5]. Indeed, some experiments would be difficult, if not impossible, without the use of general anesthesia, which suppresses awake behavioral movements and permits surgical access to the gastrointestinal (GI) tract. Despite some research on this issue [6], it remains unclear what effects general anesthesia has on these GI myoelectric signals.

Research suggests that general anesthesia inhibits the enteric nervous system and inhibits GI slow waves [7]. The slow wave is the most studied GI myoelectric signal and is believed to originate from the interstitial cells of Cajal (ICC), a component of the enteric nervous system [8]. Although an ICC coupled network is found in all parts of the GI tract, those ICC in the proximal gastric corpus are believed to initiate the slow wave contraction, mainly due to their higher intrinsic frequency [9]. There are also connections between vagal afferent and efferent fibers and the ICC network and anesthesia has been reported to affect both of these pathways [10]. Moreover, at least one study reported that general anesthesia suppresses GI slow wave responses [7].

The current study was designed to determine the effects of isoflurane anesthesia on gastric myoelectric signals. The ferret was used in this study because it is a well-known model for emetic testing and gastrointestinal measures [3,11–13], and importantly, nausea and vomiting are cardinal features of gastroparesis [14]. In this report, we conducted two experiments. First, we compared gastric myoelectric signals between awake and isoflurane-anesthetized conditions. Second, we determined whether gastric myoelectric signals were affected by body movements by comparing data during behavioral movement and rest episodes in awake animals.

## 2. METHODS AND MATERIALS

### 2.1. Animals

This study included adult purpose-bred influenza-free male ferrets (N=8; Mustela putorius furo; Marshall BioResources, North Rose, NY, USA). Table 1 provides information about the animals used in the study. Animals were adapted to the housing facility for at least 7 days before surgery. Home cage housing (62 × 74 × 46 cm) was under a 12 h standard light cycle (lights on at 0700 h), in a temperature (20–24 °C) and humidity (30–70%) controlled environment. Food (ferret kibble; Mazuri Exotic Animal Nutrition, St. Louis, MO) and drinking water were freely available. At the end of the study, euthanasia was performed with an intracardiac injection of euthanasia solution (390 mg/ml sodium pentobarbital; SomnaSol EUTHANASIA-III Solution, Covetrus, Dublin, Ohio, USA) under isoflurane general anesthesia (5%). The University of Pittsburgh Institutional Animal Care and Use Committee (IACUC) approved all experimental procedures.

**Table 1.** Subjects: Body weights (kg) and ages (days) at the start and end of the experiments, and experiment participation of each animal.

| Animal | Body weight (kg) |  | Age (days) |  | Experiment |  |
| --- | --- | --- | --- | --- | --- | --- |
|  | Start | End | Start | End | *anesthesia* | *movement* |
| 58-21 | 1.30 | 1.59 | 139 | 177 |  | x |
| **59-21** | 1.48 | 1.77 | 139 | 176 | x | x |
| **60-21** | 1.30 | 1.59 | 139 | 177 | x | x |
| 69-19 | 1.7 | 1.7 | 197 | 204 |  | x |
| **63-21** | 1.43 | 1.65 | 139 | 176 |  | x |
| **85-21** | 1.4 | 1.4 | 163 | 163 | x | x |
| **87-21** | 1.8 | 1.8 | 161 | 161 | x |  |
| **89-21** | 1.8 | 1.8 | 167 | 167 | x |  |

### 2.2. Electrode implantation surgery

Using our published procedures [3], ferrets were implanted with GI serosal surface electrodes (Fig. 1A) at four locations along the stomach axis (Fig. 1B). Anesthesia was induced by intramuscular injection of ketamine (20 mg/kg), surgical sites were shaved, and animals were endotracheally intubated. Surgery was conducted aseptically under general anesthesia using isoflurane (1–2%) vaporized in O_2_. Body heat was maintained at 36–40°C using a heating pad. Incisions on the dorsal neck and abdominal surface were used to advance a trocar tube to route the electrode leads from the skull to the abdominal cavity. Animals were implanted with four gastric electrodes (Fig. 1B), two intestinal electrodes, and one abdominal vagus nerve cuff electrode. Data collected from the intestinal and vagus nerve electrodes are not reported here. Each gastric planar electrode was attached to the GI serosal surface using 8-0 surgical silk with eight suture locations, four around each contact point. Before closing, the abdominal cavity was flushed with Cefazolin (1g/L), an antibiotic. The abdominal muscle was closed with 4-0 resorbable suture material and the skin with 3-0 monofilament nylon suture material followed by applying surgical glue to the incision site. The ferret was then turned with dorsal surface up and the electrode connectors were anchored to the skull using surgical cement. The impedances of all connections were tested (Nano2 front end, Ripple grapevine, Salt Lake City, UT) prior to the end of the surgical session. Post-surgical analgesia was provided twice a day by administering buprenorphine (0.05 mg/kg, sc) and ketoprofen (2 mg/kg, sc). Amoxicillin, anti-biotic, was administered twice a day for 10 days (20 mg/kg, po). At 14 days post-implantation, under isoflurane (2 to 4%) administered through a facemask, sutures were removed. Animals were also acclimated for three days to the test chambers prior to testing.

**FIGURE 1.**
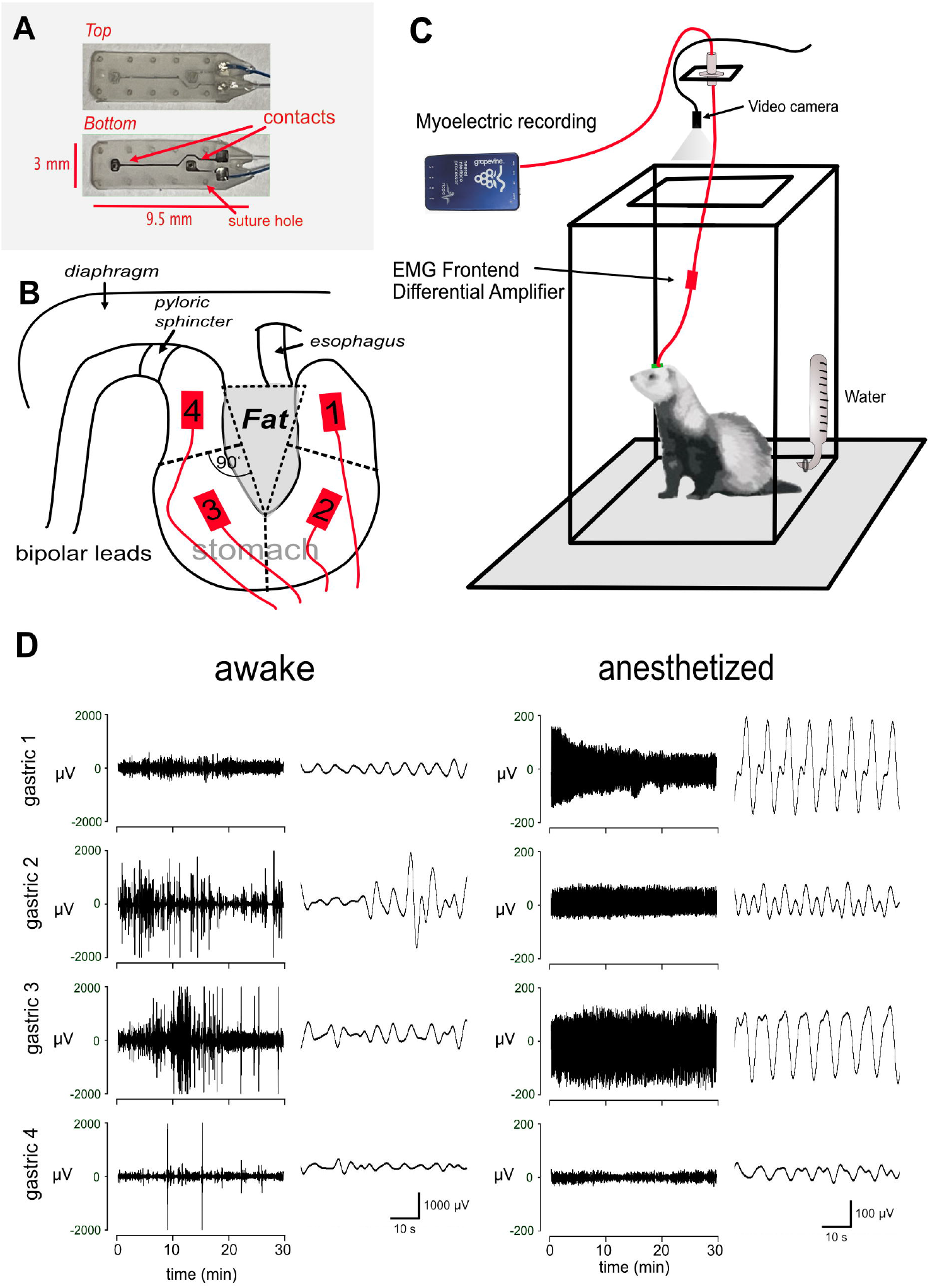
Surgical placement of GI electrodes, test chamber, and examples of gastric myoelectric data. A) GI planar electrodes. B) Placement of electrodes on the four quadrants of the stomach; reference lines are in red relative to the fat pad in the lesser curvature. C) Testing chamber with a tethered electrophysiological acquisition system to record GI myoelectric responses and video of ferret behavior. D) Examples of gastric myoelectric responses from the four channels in awake and isoflurane anesthetized conditions; data traces show all 30 min of recording and a zoomed in segment showing the gastric slow wave activity during 1 min. Note: the awake and anesthetized plots are on different voltage scales.

### 2.3. Testing

Following three hours of food deprivation to empty the stomach, animals were transported to the test room, connected to the recording tether, and placed in the behavioral test chamber (Fig. 1C). Cameras were positioned above the chamber to collect video of animal behavior for offline analysis. Animals were allowed to adapt to the chamber for 30 min before electrophysiological recording for 1 hour. Free moving testing occurred for at least 10 days before a terminal procedure on the final day of testing. Myoelectric signals were recorded with a Ripple Grapevine Neural Interface Processor and a differential EMG headstage (Ripple LLC, Salt Lake City, UT) and sampled at 2 KHz.

On the final test day, each animal was acclimated to the chamber for 30 min, and then recordings were conducted for 30 min. The animal was then placed in an anesthesia chamber, and 5%isofluorane was delivered for approximately 5 min. Subsequently, animals were removed from the chamber and anesthesia was maintained using isoflurane (1 to 3%) delivered through a face mask. Electrophysiological signals were then recorded from the gastrointestinal electrodes for 30 min. Additionally, to monitor physiological parameters, the electrocardoiogram (EKG) was recorded by placing needle electrodes placed under the skin; signals were amplified 5 to 20K times and filtered with a Model 1700 Differential AC Amplifier (A-M Systems, Sequim, WA). Rectal body temperature was recorded using a Model TC-1000 temperature controller (CWe Inc., Ardmore, PA). Respiration signals were collected with a Cambridge Electronic Design Power 1401 data collection system and Spike 2 ver. 9 software (Cambridge Electronic Design Ltd., Cambridge, England). At the end of this test, animals were euthanized. The GI tract was then extracted for later evaluation of electrode locations.

### 2.4. Behavioral scoring

Video of ferret behavior was scored off-line using BORIS software [15]. Behaviors scored included jumping/standing, digging, locomotion, urination, defecation, grooming, head shake, touching the tether, and resting. The presence of any of prior behaviors, except resting, was characterized as behavioral movement. Resting animals were considered to have “no movement.” In the final analysis, we combined all behavioral movement into one category of “movement”.

### 2.5. Preprocessing of myoelectric signals

Myoelectric signals were filtered using a second-order digital Butterworth band-pass filter with high- and low-pass cutoffs at 0.1 Hz and 0.3 Hz (6-18 cycles per minute), respectively. The filter was applied twice, forward and backward, to remove timeshifts caused by forward filtering. A filtered signal exceeding ± 2000 mV was assumed to be artifact (i.e., outside the physiological range [3,11]) and was discarded, together with 10 points preceding and succeeding the artifact, and removed data were replaced via linear interpolation.

For the awake versus anesthesia analysis, we started with 20 paired LFP time series obtained from 4 channels in 5 animals. One channel was removed because the signal failed to be recorded in the anesthetized state. Another channel was removed because there were too many artifacts (32%) in the awake state. The remaining 18 channels were retained for analysis. Ten channels -- all the channels of two animals as well as one channel for each of the three other animals -- had less 0.0001% artifacts in each state. The remaining eight channels had less than 3.7% artifacts in the awake state. One of the corresponding anesthetized state recordings had 1.1% artifacts and the seven others had less than 0.0001%. The exact proportions can be found in Table 2. Artifacts tended to happen in the awake state, alone or in small clusters, but with equal prevalence in the movement and rest period, and were possibly caused by the triboelectric effect when the leads move under the skin or as a result of EMG contamination [16]. Even though artifacts primarily happened in the awake state, it is very unlikely their removal had an effect on the substantive results of our analysis given how seldom they occurred.

**Table 2.**
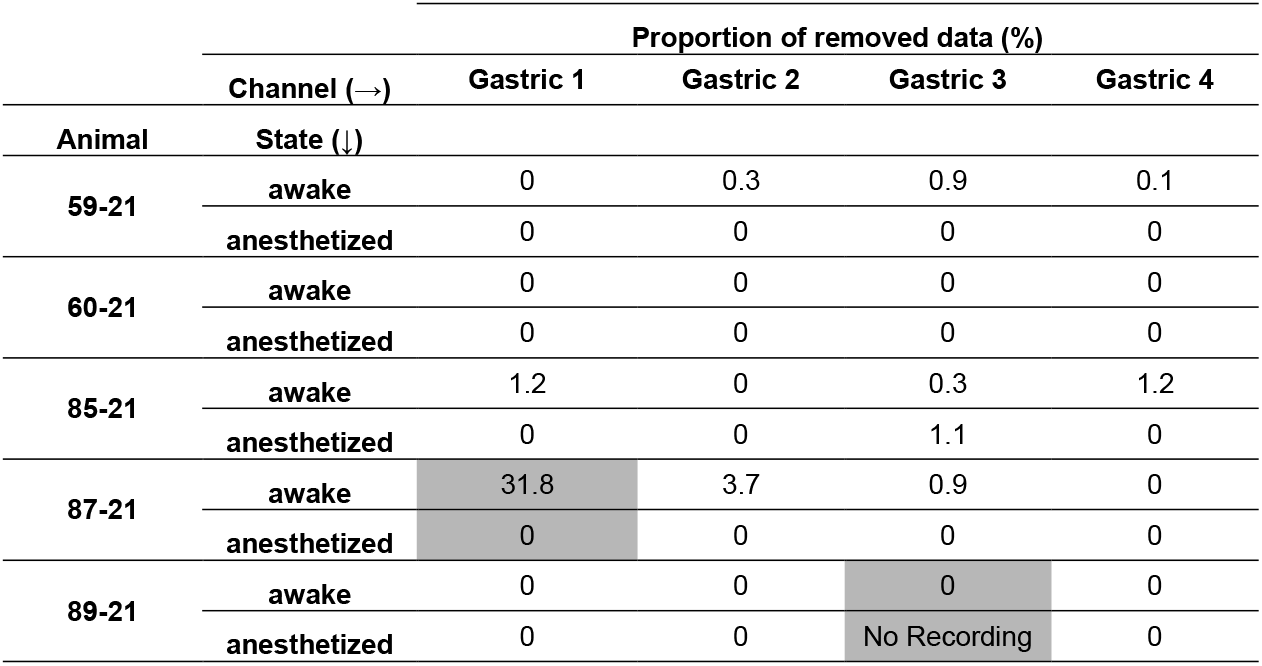
Proportion of removed data (as % of time for each state) for the anesthesia experiments due to artifact identification. Removed channels in gray.

For the movement versus no movement analysis, we retained all 28 (4 channels x 7 recording sessions) for analysis. Sixteen channels had less than 0.0001% artifacts in each state and the remaining 12 had less than 3.7%. The exact proportions can be found in Table 3. Fig. 1D shows examples of the preprocessed signals.

**Table 3.** Proportion of removed data (as % of time for each behavior) for the movement experiments due to artifact identification.

| Experiment | Channel (→)<br>Behavior (↓) | Proportion of removed data (%) |  |  |  |
| --- | --- | --- | --- | --- | --- |
|  |  | Gastric 1 | Gastric 2 | Gastric 3 | Gastric 4 |
| 85-21 | movement | 2.4 | 0 | 0.3 | 3.7 |
|  | no-movement | 0.9 | 0 | 0.4 | 0.7 |
| 69-19 B2 | movement | 0 | 0 | 0 | 0.3 |
|  | no-movement | 0 | 0 | 0 | 1.7 |
| 69-19 B3 | movement | 0 | 0 | 0.1 | 2.4 |
|  | no-movement | 0 | 0 | 0 | 0.9 |
| 58-21 | movement | 0 | 0.3 | 0.5 | 0.1 |
|  | no-movement | 0 | 0.5 | 0 | 0 |
| 59-21 | movement | 0 | 0.8 | 0 | 0 |
|  | no-movement | 0 | 1.0 | 0 | 0 |
| 60-21 | movement | 0 | 3.2 | 0 | 0 |
|  | no-movement | 0 | 1.0 | 0 | 0 |
| 63-21 | movement | 0 | 1.1 | 0 | 0 |
|  | no-movement | 0 | 0.6 | 0 | 0 |

### 2.6. Data analysis

#### Extraction of signal power and PSD with 95% confidence interval estimation

After preprocessing, we calculated signal power using a 60-second rolling Hamming window, with 48 seconds overlapping between consecutive windows. We then log-transformed the power to (i) stabilize its variability - larger power values have correspondingly larger variabilities on the original scale - and to (ii) satisfy the Gaussian assumption of the response variable (i.e., log power) in regression analyses. We estimated the power spectral density (PSD) of the signals, together with 95% confidence intervals, using Welch’s overlapped segments averaging method, with 1-minute overlapping windows [17]. For some analyses, total signal power was considered, while for others, we examined power in frequency bands around the dominant frequency (DF), namely *DF* − [3,1) *HZ* (i.e., bradygastric), *DF* ±1 *HZ* (i.e., normogastric), and *DF* +(1,3] *HZ* (i.e., tachygastric), as described previously [3].

#### Estimating the magnitude of the effect between states

We estimated the size of the effect on log power between awake and anesthetized states, and between behavioral movement and rest states, on average across all animals and channels. Letting *Y*_*ijk*_ denote log power for animal *i*, channel *j*, and state *K*, with *K*=1denoting rest (or anesthetized) and *K*=2 denoting behavioral movement (or awake), we fitted the regression model *E[Y*_*ijk]*_ =β_*ij*_ + δ_*k*_, setting δ_1_ =0. This model assumes that the mean log power for channel *j* of animal *ii* is β_*ij*_ in the rest (or anesthetized) state and β_*ij*_ + δ_2_ in the behavioral movement (or awake) state. The β_*ij*_ could be different across animals and channels but δ_2_ was kept constant, so that δδ_2_ measured the difference between the two states, averaged across animals and channels. We used a standard regression to fit this model and verified that the response variable (log power) was close to Gaussian.

## 3. RESULTS

### 3.1. Comparison of gastric myoelectric activity between awake and anesthetized states

Figure 2A shows the PSDs for all animals and all channels in the awake state and under isoflurane anesthesia. We discarded two channels (gray rectangle) due to overly noisy recordings. There is a significant reduction (*p* <.001, large-sample approximation of (1 – α)% confidence intervals from χ^2^ distribution of the PSD estimator [17]) in estimated PSDs in almost the entire frequency spectrum after exposure to anesthesia in 17 out of 18 gastric recording channels across five animals. The reduction in total power (area under the PSD curve) is over 90% (between 91.4% and 99.6%) in 13 cases, and between 34.4% and 88.3% in the remaining four cases. The normalized PSDs in Figure 2B help visualize how the power shifted in various frequencies under anesthesia compared to the awake state. We observed increased concentrations of power around the dominant frequency in 12 out of 18 channels; and the distributions of power exhibit multiple peaks in 13 out of 18 channels, whereas they are typically unimodal in the awake state.

**FIGURE 2.**
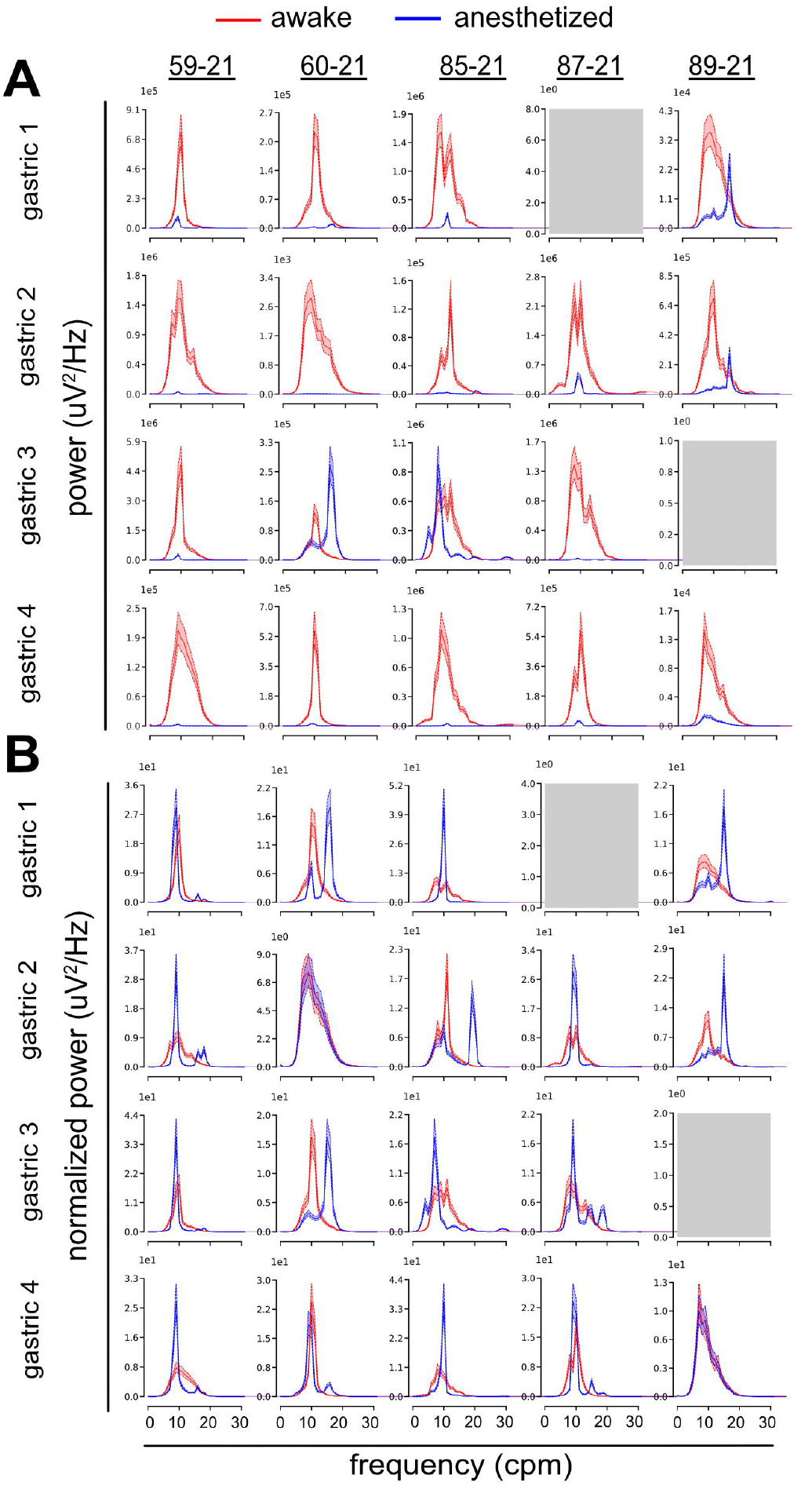
Effect of isoflurane anesthesia on power (A) and normalized power (B) of gastric myoelectric responses in each ferret. Grid of Welch’s overlapped segments averaging power spectral densities (PSD) estimated from myoelectric recordings in awake (red) and anesthetized (blue) states, from 5 subjects (columns) at 4 channels (rows), with 95% confidence interval (shaded regions). Two channels were omitted due to noisy data (grey shading). After exposure to anesthesia, we observe a significant (*p* < 0.01) suppression of power in all but one panel (4A Gastric 3, subject 60-21). Moreover, under isoflurane anesthesia power shows an increase in concentration around the dominant frequency in 12 out of 18 panels compared to the awake state (4B).

Figure 3 focuses more specifically on the normogastric, bradygastric, and tachygastric bands rather than on the full PSDs. The figure shows the average log powers in these three bands, together with 95% confidence intervals, for awake and anesthetized states, for all channels and animals. Power decreased during anesthesia compared to the awake state for all subjects and channels (*p* < 0.05; independent t-test), except for animal 60-21, channel 3, where there was no significant difference between the two states. When combining all recordings and channels, the decreases in log power in the anesthetized state were highly significant in the normogastric (2.603, SD = 0.037, *p* < .001), bradygastric (3.990, SD = 0.051, *p* < .001), tachygastric (4.175, SD = 0.052, *p* < .001) bands, as well as when power was aggregated across all three bands (2.725, SD = 0.037, *p* < .001). These numbers correspond to decreases in power of 92%, 98%, 98.5%, and 93%, respectively. The relatively less pronounced drop in power in the normogastric band is in line with the observed increase in concentration of power around the dominant frequency in the anesthetized state (Figure 2B).

**FIGURE 3.**
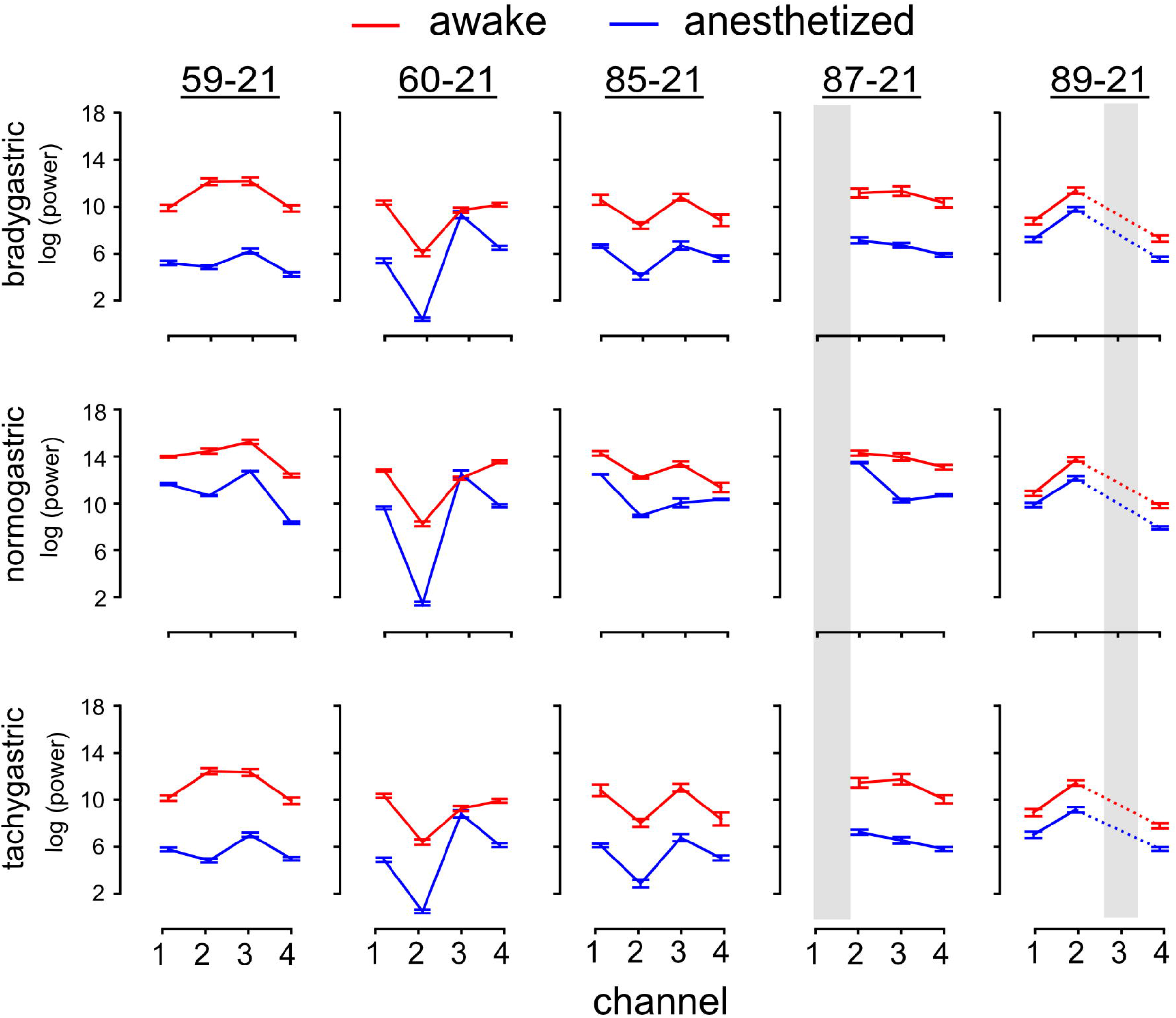
Average and 95% confidence intervals for power (log-transformed) in the bradygastric, normogastric and tachygastric bands for each animal, extracted from 30 min recording periods during the awake (red) and anesthetized (blue) states. Data that exceeded a threshold were deemed corrupted and removed (grey shaded areas); see section 2.5. In most recordings, we observe significantly greater power in the awake state (17 out of 18 channels in bradygastric, normogastric and tachygastric bands *p* < 0.05; independent t-test). Differences in power between the two states tend to be similar across the three frequency bands for each channel and subject.

### 3.2. Comparison of gastric myoelectric activity between behavioral movement and rest

Animals exhibited two characteristic types of behaviors: (i) sustained periods of either behavioral movement or rest or (ii) frequent transitions between short periods of behavioral movement and rest. Altogether, animals exhibited type (i) behavior in three recording sessions and type (ii) in four sessions. Figures 4A and 4C show the log power in the brady-, normo-, and tachygastric frequency bands for each of the four recording channels, for representative recordings of types (i) and (ii), respectively. Figure 4A suggests that for all channels, the log powers in the bradygastric and tachygastric bands are markedly reduced during rest compared to behavioral activity, and somewhat reduced in the normogastric band. The log powers decrease gradually for about 5 mins (time stamps 50 to 75 min) at the start of the second rest period and then remain around the same level (within random variations). Perhaps for this reason, it is difficult to see the same effect in Figure 4C because no rest or activity period exceeded 5 mins for recordings of type (ii). To further explore the effects of behavioral movement during type (ii) recordings, for each channel and each frequency band, we collected the log power values at all time points and gave each measurement the label “rest” or “behavioral movement”. Then we overlaid the histograms of the “rest” and “behavioral movement” values and added averages and 95% confidence intervals. Figure 4B clearly shows that for all combinations of channels and frequency bands but one (Gastric 3 - normogastric), log power is lower in rest periods compared to behavioral movement periods. The same effect also appears in Figure 4D, although to a lesser degree.

**FIGURE 4.**
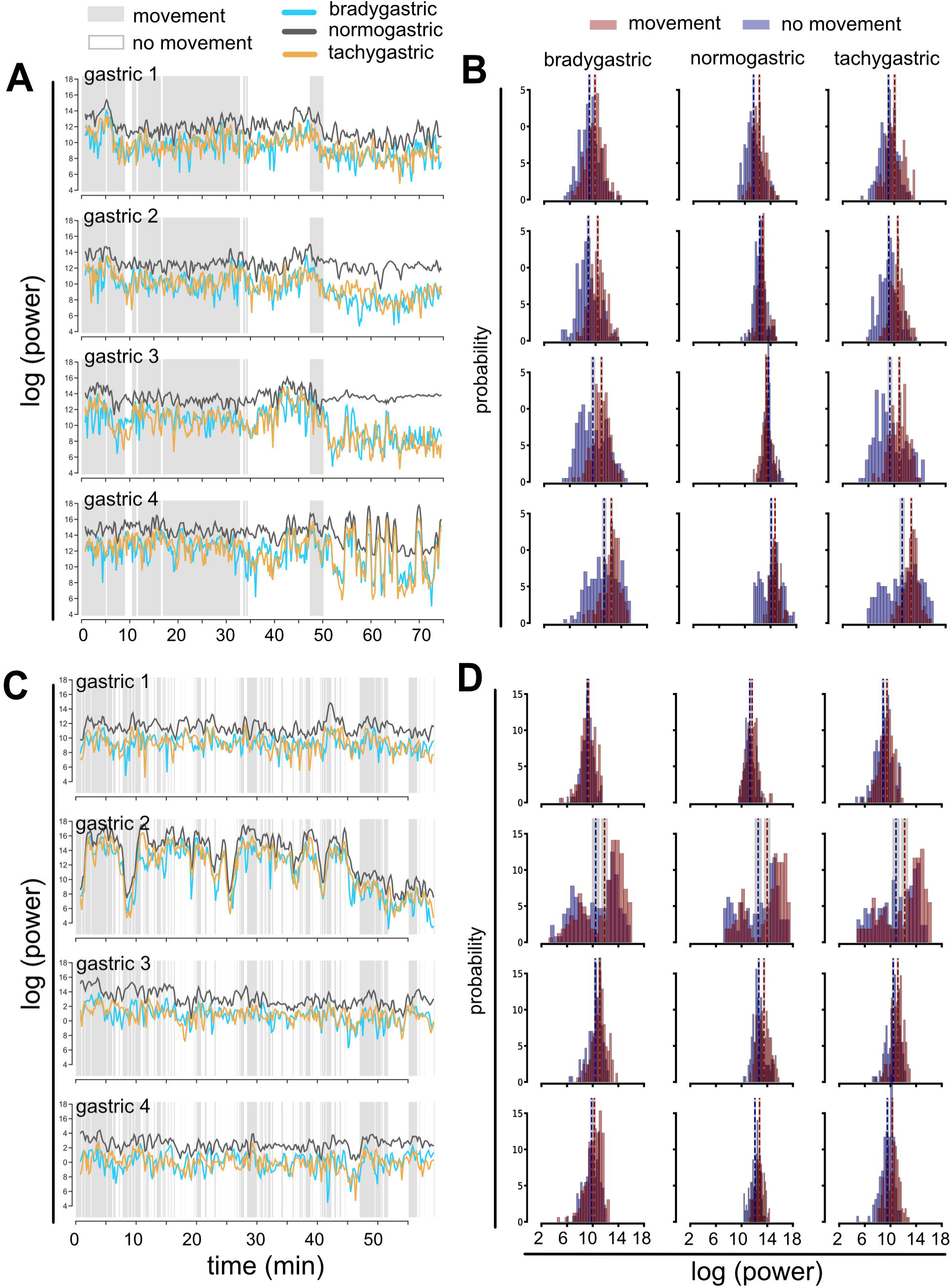
Time courses of power (log-transformed) in the bradygastric, normogastric and tachygastric bands for two representative behavioral movement sessions. (A) The subject’s behavior was characterized by sustained periods of behavioral movement or rest. (C) The subject exhibited short and interspersed periods of behavioral movement and rest. In (A), the decrease in power is evident during the prolonged rest period (from 50 to 75 min), especially for the brady- and tachygastric bands. The absence of sustained rest periods in (C) may explain why we do not observe a similar decrease. Panel sets B and D show the probability distributions of log power at each time point from panel A and C, respectively, during movement (red) and rest (blue). Dotted lines and shading regions indicate means and 95% confidence intervals. (B) The average log power is significantly greater during movement compared to rest for all combinations of channels and frequency bands (*p* < 0.05; independent t-test) except gastric 3 in the normogastric band. (D) The effect is present but smaller in 8 out of 12 channel-band pairs, *p* < 0.05; independent t-test).

Figure 5 summarizes the information in Figures 4B and 4D for all recordings. Specifically, for each recording, channel, and frequency band, the mean log power values and 95% confidence intervals are plotted for rest and behavioral movement periods. For the three recordings of type (i) (shaded region in Figure 5) the mean log powers in all bands and channels are significantly lower in rest periods compared to behavioral movement periods, albeit with a smaller, or sometimes insignificant, effect in the normogastric band. The effect can also be observed in the four recordings of type (ii), although it is smaller and more sporadic, perhaps because the interspersed periods of rest and activity are too short for the power to transition from one state to the other. The decreases in log power in rest periods compared to behavioral movement periods, averaged across all recordings and channels, are 1.002 (SD = 0.041, *p* < .001) in the bradygastric band, 1.150 (SD = 0.042, *p* < .001) in the tachygastric band and 0.628 (SD = 0.029, *p* < .001) in the normogastric band, corresponding to 63%, 68% and 47% decreases in power, respectively. Combining the three power bands, the overall decrease is 0.670 (SD = 0.029, *p* < .001) on the log scale, corresponding to a 49% decrease in power.

**FIGURE 5.**
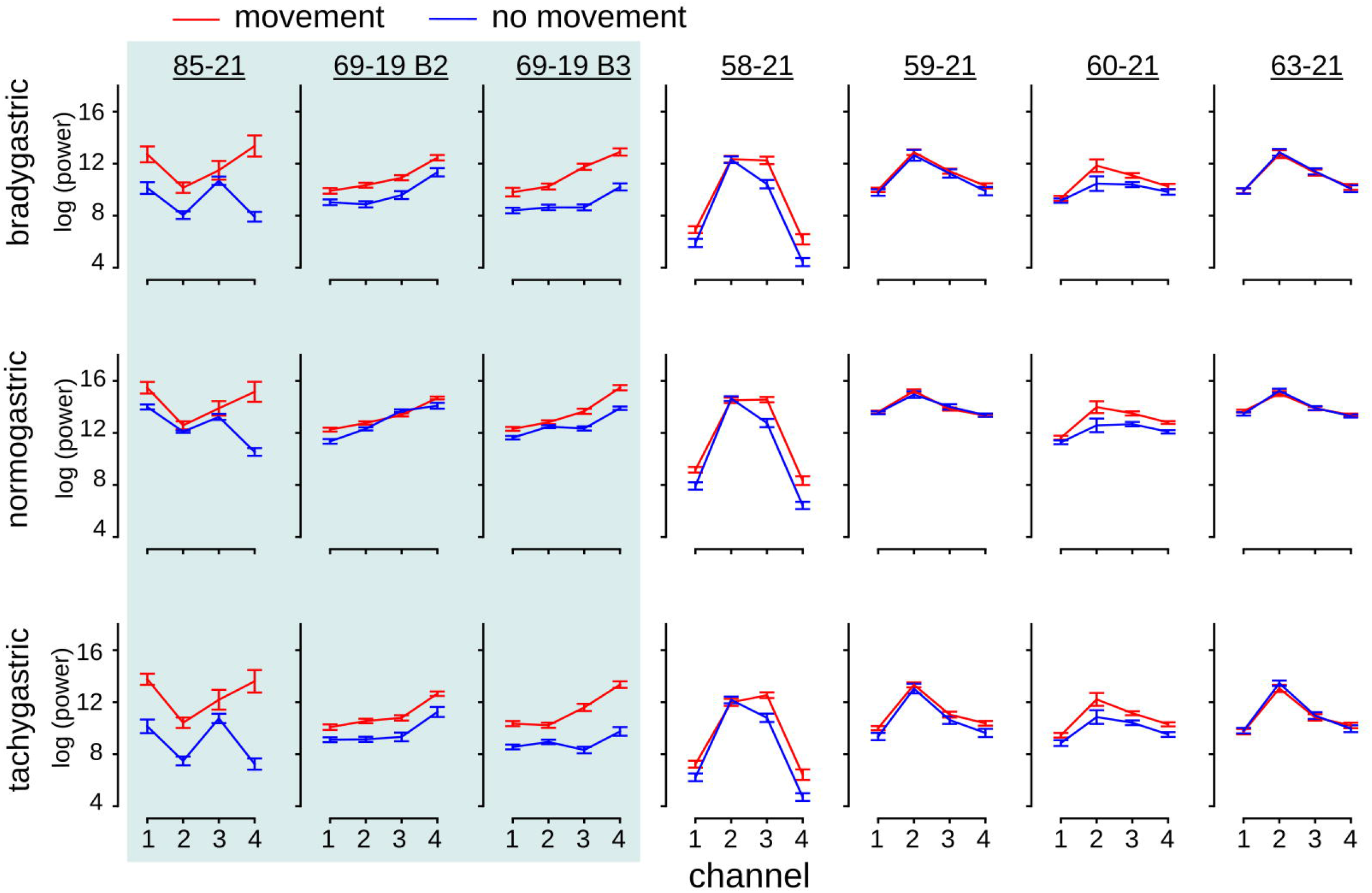
Average and 95% confidence intervals for power (log-transformed) for each behavioral movement session during movement (red) and rest (blue), broken down by frequency bands (rows) and channels. The shaded region indicates sessions characterized by sustained periods of movement and rest (type (i) sessions). In these types of recordings, the power is significantly greater during movement periods compared to rest in the bradygastric (11 out of 12 channels *p* < 0.05; independent t-test) and tachygastric bands (12 out of 12 channels *p* < 0.05; independent t-test), with a smaller effect in the normogastric band (8 out of 12 channels, *p* < 0.05; independent t-test). In the remaining four sessions of short and interspersed periods of movement and rest, the effect is present but smaller (5 out of 16 channels in bradygastric range *p* < 0.05; 8 out of 16 in normogastric range *p* < 0.05; 9 out of 16 in tachygastric range *p* < 0.05; independent t-test).

## 4. DISCUSSION

The results show that isoflurane anesthesia reduced overall signal power and increased the concentration of power around the dominant frequency of gastric slow waves compared to the awake condition. Part of these differences could be due to behavioral movements of the animal; however, the effect of behavioral movement on signal power was smaller compared to changes observed with isoflurane anesthesia.

Importantly, the current study, as far as we are aware, is the first to directly compare gastric slow wave changes between individual animals between awake and general anesthesia conditions. There is a report of the effects of general anesthesia (thiopentone, nitrous oxide, and isoflurane) compared to the awake condition in human children but the anesthesia condition also includes circumcision surgery; therefore, the effect of anesthesia alone cannot be determined [18]. The only prior study of the effects of anesthesia (thiopental, isoflurane, and nitrous oxide), without the confound of a surgical procedure, on gastric slow wave activity, was conducted in the porcine model, but did not include a direct comparison to the awake condition [6].

While there was a stark change in signal power between the anesthetized and awake states, it is important to acknowledge that animal behavioral movement itself could have contributed to the observed difference in signal power between the awake and isoflurane anesthesia conditions. The effect of behavioral movement could be the result of recording artifacts or potentially changes in myoelectric activity due to repositioning of the GI tract during movement. Since gastrointestinal function is controlled by the autonomic nervous system [19], it is also possible that the observed changes in myoelectric activity could be partially attributed to changes in sympathetic versus parasympathetic activity to the GI tract during movement.

In summary, although we observed gastric myoelectric signal power decreases during isoflurane anesthesia, we found that the pattern of slow waves was unchanged. The comparison to the awake condition, however, does not capture the variability of the slow wave signals that might occur during the daily cycles and during ingestion of water and food, which cannot be assessed in experiments under anesthesia. Future studies should address these additional variables, which could impact the utility of gastric myoelectric recordings for measuring GI responses in different states and application to patients suffering from GI motility diseases.

## AUTHOR CONTRIBUTIONS

CH, LF, BY designed the experiments. SF, MS, CH collected data. LT, MS, VV, CH, LF conducted data analysis and interpretation. CH, LT, VV drafted the manuscript. All authors approved the revisions and final version of the manuscript.

## Acknowledgements

This research was supported by funding from the NIH awards R01DK121703 and U18TR002205 from the SPARC Common Fund Program.

